# ATG16L1 WD domain regulates lipid trafficking to maintain plasma membrane integrity to limit influenza virus infection

**DOI:** 10.1101/2024.12.09.627529

**Authors:** Benjamin Bone, Luke Griffith, Matthew Jefferson, Yohei Yamauchi, Thomas Wileman, Penny P. Powell

**Affiliations:** Biomedical Research Centre, Norwich Medical School, University of East Anglia, Norwich, Norfolk NR4 7TJ, UK; Molecular Medicine Laboratory, Institute of Pharmaceutical Sciences, D-CHAB, ETH Zurich, 8093 Zurich, Switzerland and Department of Virology, Graduate School of Medicine, Nagoya University, 466-8550, Japan

**Keywords:** ATG16L1 WD domain, cholesterol, cytokine storm, influenza virus, innate immunity, interferon, interferon-stimulated gene, non-canonical autophagy, LAP, CASM, LANDO

## Abstract

The non-canonical functions of autophagy protein ATG16L1 are dependent on a C-terminal WD domain. Recent studies show that the WD domain is required for conjugation of LC3 to single membranes during endocytosis and phagocytosis, where it is thought to promote fusion with lysosomes. Studies in cells lacking the WD domain suggest additional roles in the regulation of cytokine receptor recycling and plasma membrane repair. The WD domain also protects mice against lethal influenza virus *in vivo*. Here, analysis of mice lacking the WD domain (ΔWD) shows enrichment of cholesterol in brain tissue suggesting a role for the WD domain in cholesterol transport. Brain tissue and cells from ΔWD mice showed reduced cholesterol and phosphatidylserine (PS) in the plasma membrane. Cells from ΔWD mice also showed an intracellular accumulation of cholesterol predominantly in late endosomes in cell culture. Infection studies using IAV suggest that the loss of cholesterol and PS from the plasma membrane in cells from ΔWD mice results in increased endocytosis and nuclear delivery of IAV, as well as increased IFNβ and ISG expression. Upregulation of IL-6, IFNβ and ISG mRNA were observed in *ex vivo* precision cut lung slices from ΔWD mice both at rest and in response to IAV infection. Overall, we present evidence that regulation of lipid transport by the WD domain of ATG16L1 may have downstream implications in attenuating viral infection and limiting lethal cytokine signaling.

## Introduction

Despite the importance of ATG16L1 as a member of the ATG16L1-ATG5-ATG12 complex during autophagy, where it facilitates conjugation of ATG8 mammalian orthologs such as LC3 onto double-membraned autophagosomes (1), considerable evidence points towards a number of important non-canonical functions for the protein. ATG16L1 is involved in conjugating LC3 to single-membraned compartments, such as phagosomes, during LC3-associated phagocytosis (LAP) and to endo-lysosome membranes during CASM (conjugation of ATG8 to single membranes) and LANDO (LC3-associated endocytosis) (2–4). Structural and functional analysis of ATG16L1 have identified distinct domains within the protein that contribute to these different functions (5, 6). The N-terminal ATG5-binding domain and coiled-coil domain (CCD), are required for autophagy where the presence of a WIPI binding site anchors ATG16L1 to sites of phagophore expansion (7, 8). The C-terminal WD40 domain containing seven WD repeats is not required for autophagy (9, 10), but is required for the conjugation of LC3 onto single membranes through a pathway called CASM (conjugation of ATG8 to single membranes) (7, 10). In the best characterized example of CASM an increased pH in endo-lysosome compartments activates the assembly of the vacuolar ATPase (V-ATPase) where it provides a binding site for the WD domain of ATG16L1 and subsequent conjugation of LC3. This has been called the V-ATPase-ATG16L1 axis and it can be disrupted by a Salmonella effector protein called SopF to promote replication (11, 12, 13).

ATG16L1 also contains is an extended, unstable linker region between N and C domains, which is the site of the T300A variant, a mutation associated with increased risk of Crohn’s disease and abnormalities in Paneth cell lysozyme distribution and in goblet cell morphology. We have shown previously that mice lacking the linker and WD domain of ATG16L1 (ΔWD) up to glutamate at position 230 develop normally and maintain tissue homeostasis ‘*in vivo’* (7). LC3II production and degradation of p62 in MEFs and tissues from the ΔWD mice are the same as in littermate controls showing that the ATG16L1 N-terminal domain expressed in these mice is sufficient to maintain canonical autophagy. In contrast, bone marrow derived macrophages from ΔWD mice show defects in LC3 associated phagocytosis (LAP) and antigen presentation (10, 14, 15). Studies using ΔWD mice also show that the WD domain regulates cytokine receptor trafficking during IL-10 signaling (16) and is required for recycling beta-amyloid receptors in primary microglia which is essential for maintaining cognitive health, with mice lacking the WD domain developing spontaneous Alzheimer’s disease (17). We have also shown that the WD domain also plays an important role in protecting mice from influenza virus A (IAV) infection (14). These ΔWD mice are highly susceptible to IAV infection, with increased weight loss, virus lung titer, and mortality following IAV infection when compared to WT littermate controls and increased virus replication in the airways leads to a cytokine storm, pneumonia and increased mortality, thus raising the possibility of a yet undescribed role for this domain that may directly or indirectly provide antiviral protection. While decreased degradation by the LAP/CASM/LANDO pathways due to dysfunctional conjugation machinery may explain the heightened susceptibility, at present direct evidence of the WD domain and linker region in virus degradation is lacking. Cell culture experiments have shown that this region slows endocytosis of IAV and fusion of IAV envelope with endosomes resulting in slower delivery of IAV genomes into cells and delayed cytokine signaling (14). Biophysical characteristics of membranes, such as stability, are reliant upon the cholesterol concentration, with it being widely understood that the cholesterol concentrations of interacting membranes can influence viral infection (18), including IAV infection (19,20). An example is the antiviral actions of interferon-inducible transmembrane protein 3 (IFITM3) which disrupts cholesterol homeostasis to raise endosome cholesterol to slow virus entry (21). Taken together, in this study we hypothesize that the WD domain may regulate cellular cholesterol homeostasis to slow viral entry. Indeed, recent data has demonstrated that ATG16L1 is involved in lysosomal exocytosis, which promotes plasma membrane repair following membrane damage by bacterial pore-forming toxins, and during Listeria monocytogenes infection, a function which limits cell to cell spread (22). Plasma membrane repair involves efflux of cholesterol from lysosomes to the plasma membrane, a process dependent upon the WD domain of ATG16L1 and the ATG5-ATG12 conjugant, but not reliant upon other proteins crucial for autophagy (23).

We build on our previous study (14) and find an unconventional role for the WD domain in protecting mice from IAV infection through control of cholesterol distribution. We performed *ex vivo* infection challenges of precision cut lung slices from WT and ΔWD mice and show increased infection and elevated cytokine expression in ΔWD tissue. Following this, we monitored by fluorescence microscopy and qPCR the early infection events of IAV, observing a clear enhancement IAV entry into in ΔWD primary mouse embryonic fibroblasts (MEFs). To investigate whether cholesterol has a role in this viral entry, we used probes and chemical assays for cholesterol to show that there was an altered intracellular distribution with accumulation in late endosomes in ΔWD cells, and a depletion in the plasma membrane in brain tissue in ΔWD mice. Pharmacological modification of cholesterol distribution restores cholesterol and phosphatidylserine to plasma membrane and impacts IAV infectivity in ΔWD cells and tissues. Taken together, we show that the WD domain of ATG16L1 maintains cholesterol in the plasma membrane *in vivo*, and we argue that this slows IAV escape from endosomes and attenuates the innate immune response to IAV. This work adds to the number of unconventional activities of the WD domain of ATG16L1.

## Results

### ATG16L1 WD domain suppresses cytokine production by slowing IAV endocytosis

To build on the findings from our previous study which compared IAV infection of ΔWD and WT mice (14), we extended our work to investigate IAV infection in a precision cut lung slice tissue *ex vivo* model comprising epithelial, mesenchymal and resident immune cell types. Lung slices from WT and ΔWD mice (Fig 1A) were infected with IAV X31 for 2 hours before infection media was removed and slices were incubated in media for 24, 48 and 72 hours post infection (hpi). Virus secreted into media over 24 to 72 hpi was titrated on MDCK cells by plaque assay (Fig 1B). As previously seen, virus production was higher at all time points in lung slices from ΔWD mice compared to WT controls, this difference reaching statistical significance at 48 and 72 hours post infection (Fig 1B). Expression of mRNA for cytokines ISG15, IFITM3 and IFIT1 and the proinflammatory cytokine IL-6 was analyzed, with elevated levels observed in ΔWD lung slices compared to WT controls at 16 hpi (Fig 1C), the average relative quantity of IFITM3 and IL-6 increased nearly 3-4-fold. The results confirm our previous work and establish that the WD domain of ATG16L1 reduces IAV replication in lung slices *ex vivo* and that this reduces inflammation by reducing interferon signaling and pro-inflammatory cytokine production. ln this way the lung slices taken from ΔWD mice recapitulate the cytokine storm induced by IAV in the ΔWD mice *in vivo* (14). To analyze IAV entry into MEFs early in infection, we performed *in vitro* assessments of viral entry to confirm our previous work that the WD domain of ATG16L1 slows endocytosis and nuclear entry of IAV (14). We used two antibodies to distinguish between internal (HA1 Ab) and external virus (PINDA Ab). External HA epitopes were masked using PINDA, and then cells were permeabilized and internalized virus was identified using HA1 Ab (24). There was an increased frequency of internalized particles (green only puncta) in ΔWD MEFs cells (Fig 2A), indicating increased endocytosis of IAV into ΔWD MEFs at 30 minutes. WGA stained both the cell membrane and the nucleus after fixation. No detectable puncta were observed in cells not infected with IAV but subjected to the staining procedure. A control using pretreatment with Dynasore, a non-competitive reversible inhibitor of dynamin that inhibits endocytosis showed a distinct lack of green puncta, signifying internalized virus (Fig 2A). This suggested the WD domain has a role that provides a protective benefit to the cell membrane against viral entry by endocytosis.

**Figure 1.**
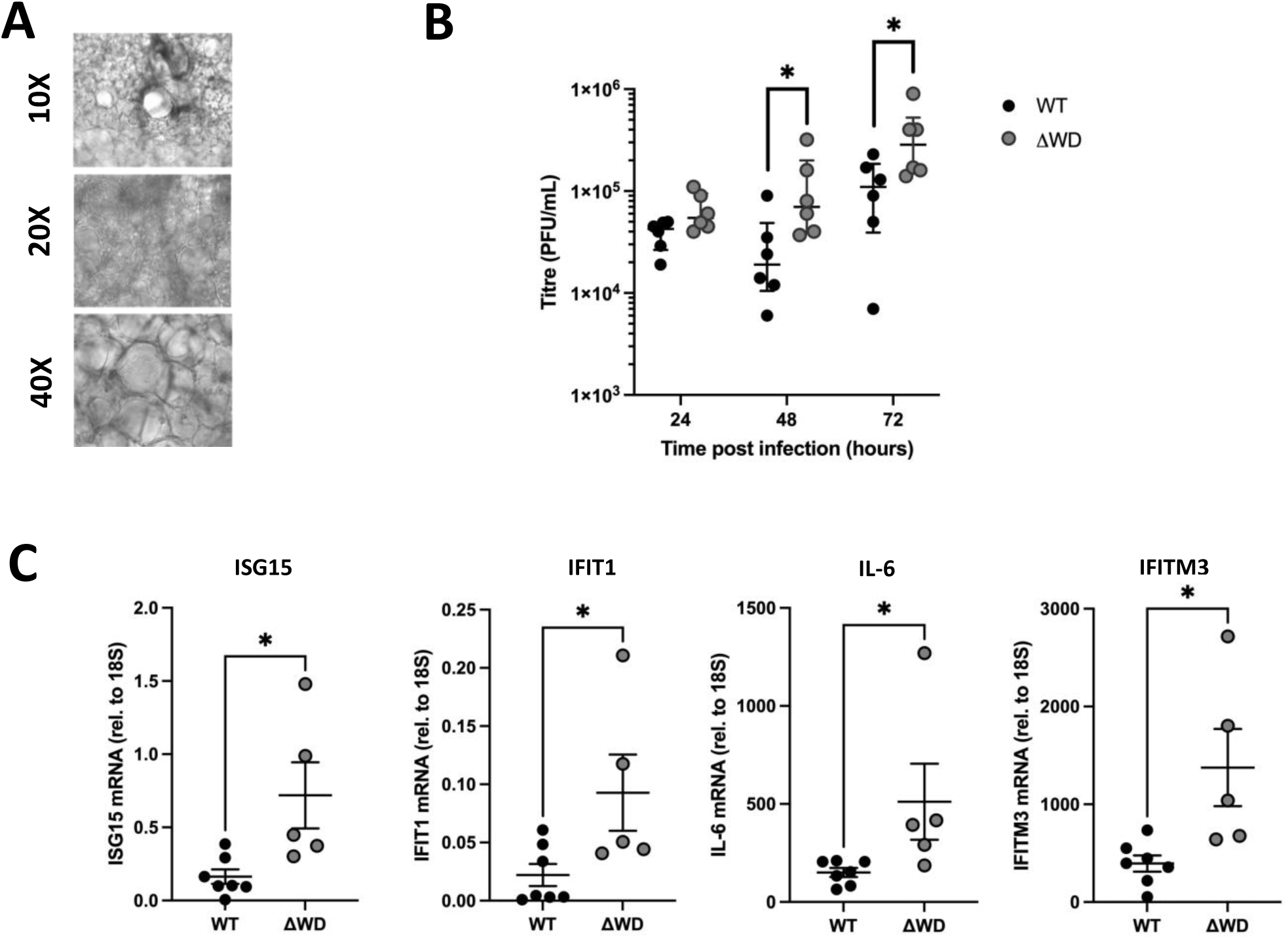
Lung explants from ΔWD mice demonstrated increased IAV replication and cytokine responses. (**A**) *In vivo* brightfield images of mouse lung explants at 10x, 20x and 40x magnification showing lung architecture of vessels, bronchioles, and cilia. (**B**) Titration of IAV X31 replication in WT and ΔWD lung slice tissue independent slices from 6 mice, at 24, 48 and 72 hpi, Virus was titrated on MDCK cells and PFU/mL (+SEM) shown per slice for the different time points. Mann Whitney U test: * = p<0.05. n=6. (**C**) Relative expression (+SEM) of IFIT1, ISG15, IFITM3 and IL-6 mRNA, normalized to 18S, at 16 hours post IAV challenge in WT and ΔWD lung slices. Mann Whitney U test: * p<0.05 n=7 WT, n=5 ΔWD.

**Figure 2.**
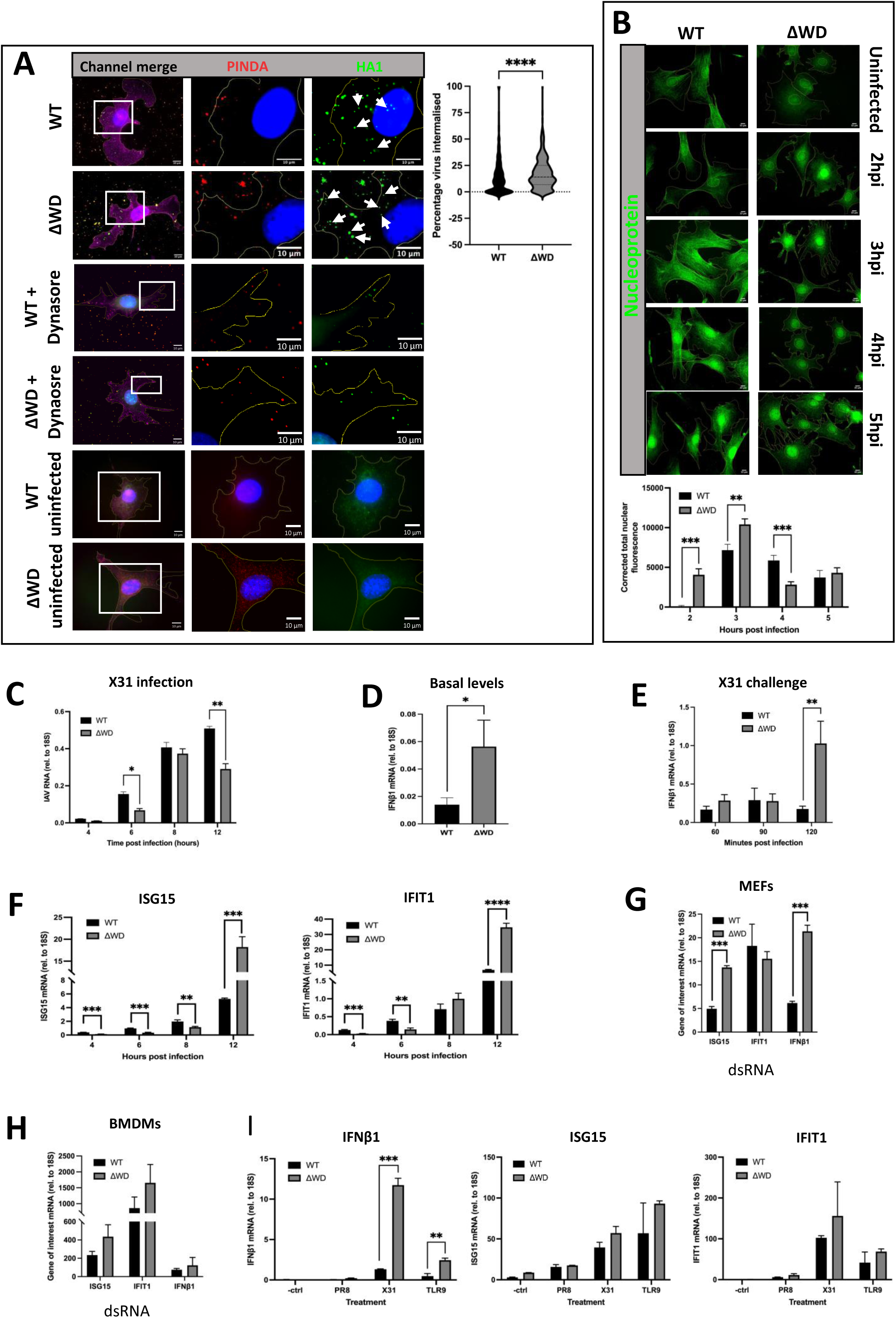
ATG16L1 WD domain slowed IAV entry and attenuates cytokine response at homeostasis and following IAV and dsRNA challenge. (**A**) IAV was bound to WT and ΔWD MEF cells for 60 mins on ice before internalization at 37 C for 30 minutes. Red puncta stained with PINDA are external virus, green puncta stained with HA1 are internalized virus (white arrows) and yellow are double stained external virus which were identified by the spot detection algorithm using CellProfiler. Controls include incubation with 80 µM Dynasore, a non-competitive reversible inhibitor of dynamin, for 30 mins, and uninfected mock control. Cells were counterstained with WGA at 647nm to show plasma membrane (magenta) and DAPI (blue). Percentage of virus internalized per cell is presented as a violin plot. A Mann-Whitney U test was performed: **** = p<0.0001, 837 WT and 417 ΔWD cells were analyzed across 3 independent coverslips. (**B**) WT and ΔWD cells were infected with IAV X31 over a time course of 0-5 hpi, fixed and stained with anti-NP antibody (green). The graph shows the average corrected total nuclear fluorescence determined for each cell nuclei (+SEM). A Mann-Whitney U test was performed: ** = p<0.01 *** = p<0.001, 30 cells were analyzed across 4 independent coverslips. (**C**) Relative quantity of IAV X31 viral RNA (+SEM) in WT and ΔWD cells over a time course of 4 to 12 hpi by qPCR. Independent samples t-test performed: * = p<0.05 ** = p<0.01. n=3. (**D**) Basal expression levels of IFNβ1 mRNA (+SEM) in uninfected WT and ΔWD MEFS, measured by qPCR. Mann Whitney U test performed: * = p<0.05 n=5. (**E**) IFNβ1 mRNA expression relative to 18S after 0-120 mins time course of IAV infection in WT and ΔWD MEFs. Mann Whitney U test: ** = p<0.005. Independent samples t-test: *** = p<0.001. n=3. (**F**) ISG15 and IFIT1 mRNA expression levels (+SEM) relative to 18S after 0–12-hour infection with IAV X31 in WT and ΔWD MEFs. Independent samples t-test: ** = p<0.005. Independent samples t-test: *** = p<0.001. n=3.. (**G**) ISG15, IFIT1 and IFNβ1 mRNA expression levels relative to 18S in MEFs from WT and ΔWD treated with dsRNA for 4 hours, measured by qPCR. Independent samples t-test: *** = p<0.001. n=3. (**H**) ISG15, IFIT1 and IFNβ1 mRNA expression levels relative to 18S in BMDMs from WT and ΔWD treated with dsRNA for 4 hours, measured by qPCR. Independent samples t-test performed: no significant comparisons detected. n=3. (**I**) IFNβ1 ISG15 and IFIT1 mRNA levels relative to 18S in BMDMs infected with IAV PR8 or IAV X31 for 4 hpi, or treated with TLR9 ligand (5µM) for 4 hours. Independent samples t-test: ** = p<0.01. *** = p<0.001. n=3.

In parallel experiments IAV nucleoprotein (vRNP) import into the nucleus was monitored by immunostaining for NP (24). A semi-quantitative estimate of delivery of vRNPs to the nucleus determined by subtracting background staining demonstrated differences in time frame of nuclear localization in the two cell lines (Fig 2B). Distinct vRNP localization to the nuclei was increased in ΔWD MEFs at 2 hours compared to 4 hours for WT MEFs. Corrected total nuclear fluorescence (CTNF) calculations were markedly greater for ΔWD values compared with WT MEFs at 2 hours and 3 hours. At 2 hours NP was virtually absent from the nucleus of WT MEFs, however by 5 hpi values between WT and ΔWD were not statistically different (Fig 2B). This suggests that in the initial stages of IAV infection, entry is optimized in ΔWD MEFs compared to WT MEFs, corroborating previously published results that reported the increased entry in cells lacking the WD domain (14). However, quantitative PCR (qPCR) analysis of viral RNA over this early time course from 4-12 hpi showed that there was a clear increased replication of IAV in WT cells compared with ΔWD cells. Genome replication was evident by 6 hpi, with elevated levels of IAV RNA in WT cells than ΔWD cells for every time point up to 12 hpi (Fig 2C). However, secretion of virus from ΔWD lung explants overtook that of WT explants from 24 to 72 hpi, as measured by plaque assay (Fig 1B).

To explain these results, we measured IFNβ1 mRNA levels at homeostasis in WT and ΔWD MEFs by qPCR and showed that basal levels of IFNβ1 mRNA were raised in ΔWD MEFs, suggesting a constitutive, low-level production of IFNβ1 mRNA in the absence of the WD domain (Fig 2D). As this transcript is normally rapidly turned over, it indicates persistent signaling or decreased suppression in ΔWD cells. This raised the possibility that the WD domain of ATG16L1 suppresses IFN signaling pathways in WT cells. Assessment of IFNβ1 mRNA production following infection of WT and ΔWD MEFs with IAV showed similar induction of IFNβ1 mRNA in both cell types during the first 90 minutes, but there was a dramatic 5-fold increase in IFNβ1 transcription at 120 min in ΔWD MEFs compared to controls (Fig 2E). This rise in IFNβ1 at 2 hours may promote antiviral responses absent from WT cells and explain the decreased replication of IAV genomes seen in ΔWD compared to WT controls up to 12 hpi. We analyzed expression of interferon stimulated genes (ISGs) that are downstream of IFNβ1 signaling and that inhibit viral RNA and protein synthesis as well as enhance virus degradation. ISG15 and IFIT1 (ISG56) expression were detected 12 hours post infection with levels of ISG15 and IFIT1 mRNA 5-fold greater in ΔWD MEFs compared to WT controls (Fig 2F).

Further experiments to elucidate signaling pathways used poly I:C as a potent dsRNA mimic and TLR3 agonist (25) to investigate differences in IFN signaling between ΔWD and WT MEFs (Fig 2G) as well as innate immune cells such as bone marrow-derived macrophages (BMDMs) (Fig 2H). Responses in BMDM were 100-fold greater than MEFs. In both MEFs and BMDMs induction of IFNβ1 and ISG15 mRNA after poly I:C stimulation was greater in ΔWD cells compared to WT cells (Fig 2F), but the difference only reached statistical significance in MEFs. These results show that the WD domain of ATG16L1 suppresses IFN signaling in epithelial cells and bone marrow-derived macrophages, not only in response to viral infection, but also to dsRNA. Increased IFNβ1, ISG15, and IFIT1 mRNA expression in BMDMs from ΔWD cells was also seen with the TLR9 ligand (Fig 2I). Interestingly, IAV strain PR8 did not illicit the same cellular response as IAV X31 (Fig 2I), a finding demonstrated in a comparative study on the IAV isolates (26).

We next investigated the antiviral sensors involved in induction of interferon. TLR3 recognizes dsRNA in endosomes, and activation results in downstream induction of IFN regulatory factors (IRFs), ultimately initiating IFN, ISG and cytokine expression. When TLR3 was inhibited by treatment with a direct, competitive, and high affinity inhibitor, there was decreased poly IC-stimulated IFNβ1 mRNA and ISG15 mRNA expression in both WT and ΔWD cells (Fig S1A). RIG-I is a cytoplasmic receptor that senses RNA with 5’-tri or di phosphate at their terminus. Knock out of RIG-I expression was validated by reduced protein production, determined by Western blot, in the WT RIG-I KO #2, and ΔWD RIG-I KO #1 and #2 cell lines (Fig S1B). WT RIG-I KO #2 and ΔWD RIG-I KO #2 cells were infected with IAV for 120 min, and IFNβ1 mRNA was measured by qPCR (Fig S1C). Lack of RIG-I protein did not affect levels of IFNβ1 mRNA, in fact, it appeared to lead to an increase in IFNβ1 mRNA levels in both WT KO cells and ΔWD KO cells compared to WT and ΔWD cells where functional RIG-I was present. Interestingly, when TLR3 was inhibited, there was an attenuation in IFNβ mRNA production following IAV infection in both WT and ΔWT cells. These results suggested that the IAV-induced interferon signaling was generated through TLR3 activation in the endosome and not by RIG-I activation in the cytoplasm, indicating an increased entry of IAV by endocytosis for this immune signaling pathway to be triggered, which supports the increased endocytosis entry into ΔWD cells reported in Fig 2A.

### Exogenous cholesterol reverses enhanced entry of IAV in ΔWD cells

Given that knock out of ATG16L1 has been reported to result in intracellular cholesterol accumulation (22, 23), as well as the widely reported relationship between cholesterol, membrane rigidity and its effect on viral infection, the next experiments explored the possibility that loss of the WD domain affected cholesterol localization within the cell and increased entry of IAV. Acid bypass was used to follow the ability of IAV to fuse with the plasma membrane. Virus was incubated with cells at 4^0^ to allow binding to the plasma membrane in the absence of endocytosis. Cells were then warmed to 37°C in acidic (pH 5.0) media at pH 5.0 to induce fusion of the virus with the plasma membrane and entry of viral RNA genomes assessed through the from staining of viral NP in the nucleus. A representative image shows increased NP in the nucleus of ΔWD cells compared to control 30 minutes following acid bypass (27) in WT and ΔWD cells (Fig 3A). We observed a pronounced increase in intracellular NP frequency The frequency of nucleoprotein (NP) puncta during a 10-50 min infection at pH5following acid bypass in ΔWD MEFs (in Fig 3B) shows a significant increase in intracellular NP frequency in ΔWD MEFs. These increases were not seen in control experiments carried out at pH7 which prevents fusion with the plasma membrane (Fig 3B). The next experiment tested whether changes in cholesterol at the plasma membrane resulting from the loss of the WD domain could influence fusion of IAV. This was tested by pharmacologically altering plasma membrane cholesterol concentration. U18666A is a cationic sterol that inhibits Niemann-Pick C1 (NPC1) protein. Inhibition of NPC1 slows movement of cholesterol from late endosomes and lysosomes and depletes cholesterol at the plasma membrane (28). 25HC is an oxysterol and metabolite of cholesterol found in the plasma membrane that interferes with viral infection via multiple pathways, including viral entry by inhibiting membrane fusion (29–31). Cells were pre-incubated with either U18666A, 25HC or cholesterol, before drug containing media was removed, and IAV uptake was tested by acid fusion at pH5 (Fig 3C). Frequency of NP puncta within WT cells and ΔWD MEFs was substantially increased when cells were pre-incubated with U18666A, suggesting cholesterol depletion at the plasma membrane enhanced fusion. In contrast, addition of cholesterol and 25HC decreased NP frequency in both cell genotypes (Fig 3C). Of note, supplementation of the cells with exogenous cholesterol resulted in a lower internalized NP frequency in ΔWD MEFs, showing that the increased entry of IAV seen in ΔWD cells was reversed by adding cholesterol.

**Figure 3.**
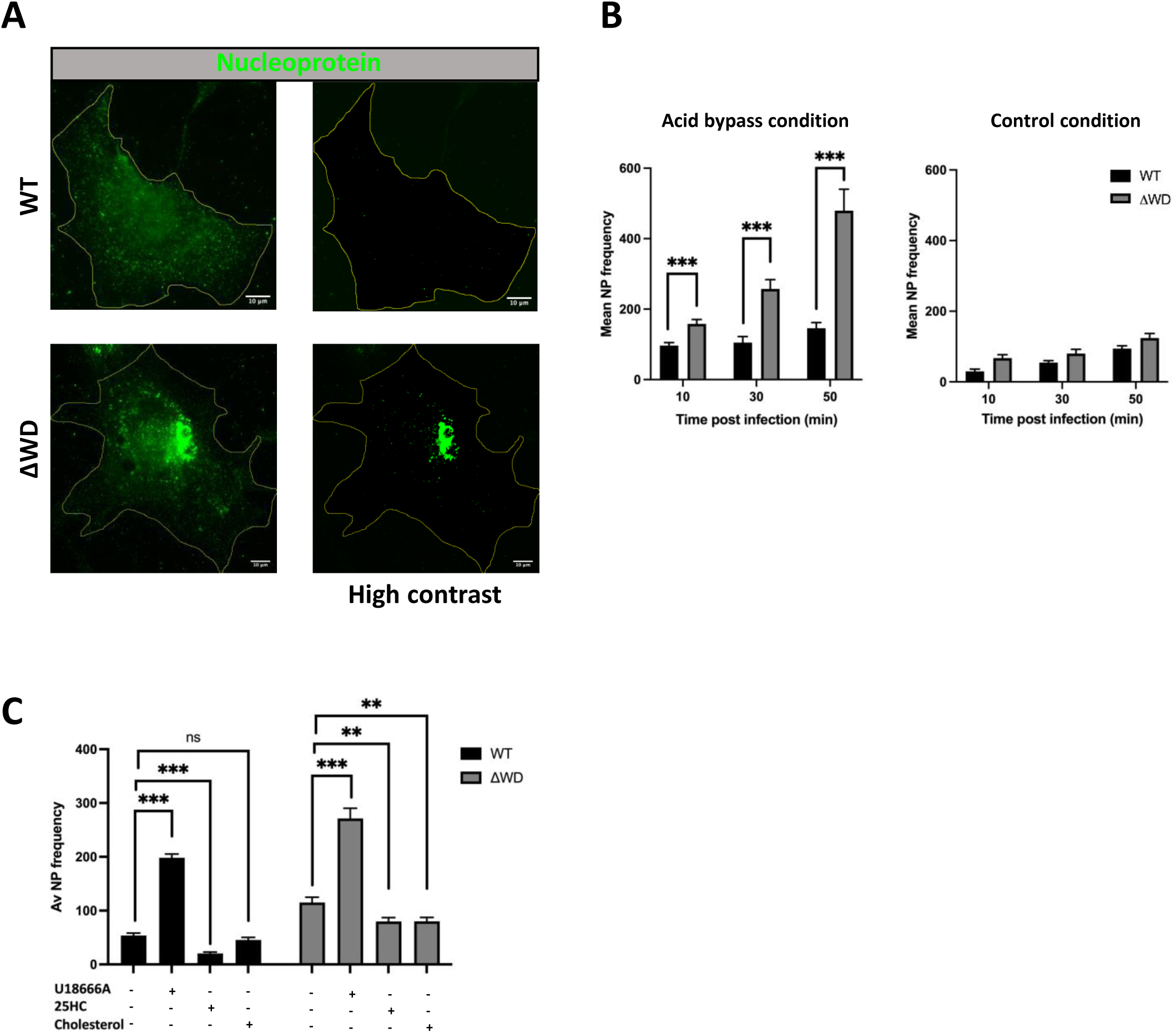
WD domain of ATG16L1 slowed fusion of IAV to the plasma membrane. (**A**) Time course of nucleoprotein puncta frequency (+SEM) in WT and ΔWD MEFs of IAV NP at 30 mins using anti-NP 488. (**B**) Graphs show a time course of frequency (+SEM) of NP puncta in WT and ΔWD MEFs at 10, 30 and 50 mins MEFs cells following acid fusion at pH5 and or control pH7. Mann Whitney U test: *** p<0.001, 50 cells analyzed across 3 independent experiments. (**C**) Nucleoprotein puncta frequency (+SEM) in WT and ΔWD MEFs cells at 50 mins post infection after pre-treatment with U18666A (3 µg/mL) for 24 hours, or 25 hydroxy cholesterol (5 µM) for 16 hours or cholesterol (80 µM) for 12 hours as indicated. Mann Whitney U test: *** p<0.001, ** p<0.01.

### Changes in intracellular cholesterol localization in cells lacking the ATG16L1 WD domain

The acid bypass experiments showed that depletion of cholesterol from the plasma membrane of WT MEFS mimicked the effects of loss of the WD domain on IAV fusion. Given that previous studies have shown that loss of the entire ATG16L1 protein leads to an accumulation of cholesterol in the cytoplasm (22), we tested whether cholesterol accumulation may also occur following loss of the WD and linker domains. Confocal microscopy on cells stained with filipin III to detect endogenous cholesterol, or on cells transfected with D4H mCherry, a plasmid expressing a modified domain 4 of perfringolysin O that effectively binds cholesterol in intracellular membranes (32, 33) showed large perinuclear puncta in the cytoplasm of ΔWD MEFs (Fig 4A). In contrast these large puncta were absent from WT cells, where filipin III and D4H cherry localized at the plasma membrane and in small cytoplasmic puncta (Fig 4A). The location of phosphatidylserine (PS), which is essential for maintaining and stabilizing cholesterol in the inner leaflet of the plasma membrane (33, 34), was detected using the PS biosensor Lact-C2-GFP (Fig 4B) and shows significantly more co-localization with cholesterol in intracellular puncta in ΔWD cells than in WT cells. This was supported by Pearson correlation analysis which determined a 3-fold increase in co localization between PS and cholesterol in ΔWD MEFs compared to WT MEFS (Fig 4B). Lact-C2-GFP localized to the plasma membrane and cytoplasmic puncta in WT cells (Fig 4C). In contrast, Lact-C2-GFP staining at the plasma membrane was noticeably reduced in ΔWD MEFs (Fig 4C). These data suggested that loss of the ATG16L1 WD domain disrupts intracellular cholesterol homeostasis leading to reduced cholesterol and PS at the plasma membrane and retention of cholesterol and PS within intracellular vesicles in ΔWD MEFs. Cells were treated with T0901317, which is an agonist of the transcription factor liver X receptor (LXR) and increases ABCA1 transporter expression (30,31).

**Figure 4.**
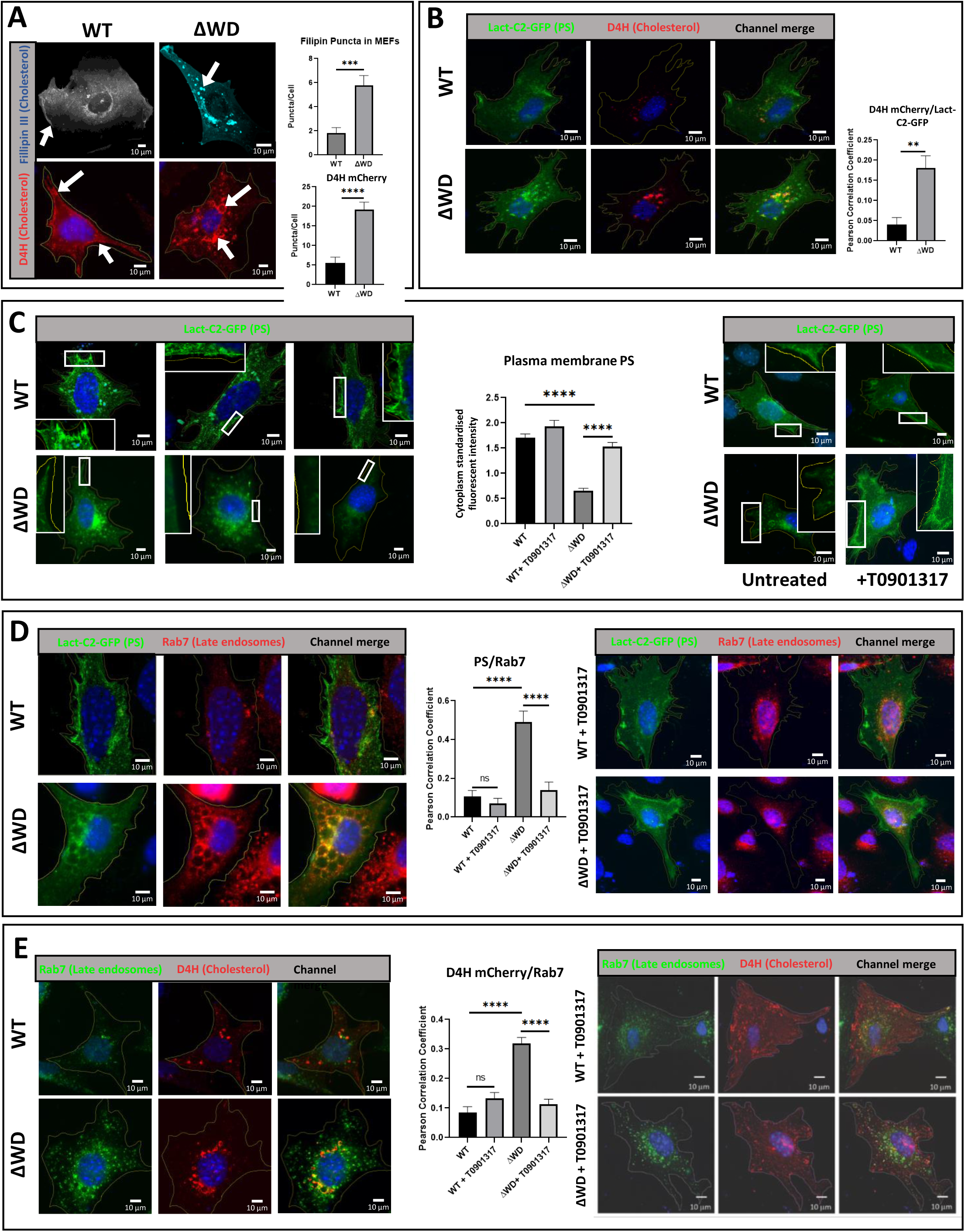
Intracellular cholesterol and phosphatidyl serine (PS) accumulated in late endosomes and lysosomes in MEFs lacking the WD domain. (**A**) Increased size and frequency of cholesterol aggregation in ΔWD MEFs. WT and ΔWD MEFs were fixed and stained with filipin III or transfected with a plasmid encoding D4H mCherry for 24 hours before fixing and staining with DAPI and viewed by Zeiss LSM980-Airyscan confocal microscope. White arrows show cholesterol at plasma membrane in WT and intracellular aggregation in ΔWD. Top graph: Filipin puncta per cell in WT and ΔWD MEFs p=0.0003, unpaired T-test with Welch’s correction, WT n=13, E230 n=17. Bottom graph: D4H mCherry puncta per cell (+SEM). Mann Whitney U test: * p<0.05, n=10. (**B**) Increased phosphatidylserine (PS) and cholesterol in cytoplasmic puncta in ΔWD MEFs. WT and ΔWD MEFs were transfected with D4H mCherry and PS biosensor Lact-C2-GFP plasmids for 24 hours before fixing and imaging. Graph shows number of double positive puncta (+SEM). Mann Whitney U test performed: ** p<0.01, n=8. (**C**) PS loss from plasma membrane in ΔWD MEFs is restored following treatment with T0901317. Untreated (left microscopy images) and T0901317 treated (right microscopy images) WT and ΔWD MEFs were transfected with the Lact-C2-GFP plasmid for 24 hours. White boxes are enlarged plasma membrane from image. Right panel shows WT and ΔWD cells treated with T0901317 for 24 hours before fixing and staining with DAPI. Cytoplasm standardized fluorescent intensity of membrane is presented on the graph (+SEM). Mann Whitney U test performed: **** = p<0.0001 N=8. (**D**) PS accumulates in large Rab7-positive cytoplasmic vesicles in ΔWD cells and restored to plasma membrane by T09011317 treatment. Untreated (left microscopy images) and T0901317 treated (right microscopy images) WT and ΔWD MEFs were transfected with the Lact-C2-GFP plasmid for 24 hours before fixing and staining with anti-Rab7 594. PS is seen with Rab7 in large vesicles of untreated ΔWD cells. T0901317 drug treatment significantly reduces ΔWD MEF PS-GFP/Rab7 puncta in ΔWD and restores plasma membrane staining. Graphs quantify PS/Rab7 col-localized puncta (+SEM). Mann Whitney U test performed: **** = P<0.0001 WT n=9, ΔWD n=8. (**E**) Intracellular cholesterol accumulation in ΔWD MEFS is reversed following treatment with T09011317. Untreated (left microscopy images) and T0901317 treated (right microscopy images) WT and ΔWD MEFs were transfected with the D4H mCherry plasmid and stained with anti-Rab7 488. Graphs quantify D4H mCherry/Rab7 col-localized puncta (+SEM). Mann Whitney U test performed: **** = p<0.0001 n=8.

Upregulation of ABCA1 on endosomal membranes facilitates active transport of cholesterol out of cells as esterified cholesterol. We show here that treatment of ΔWD cells with T0901317 restored PS to the plasma membrane (Fig 4C).

The distribution of cholesterol and PS in endosome compartments was analyzed by immunostaining for Rab7, which in its active (GTP) form binds to the cytoplasmic face of late endosomes. Analysis of WT and ΔWD MEFs using the Lact-C2-GFP probe for PS showed co-localization of PS with Rab7 in cytoplasmic puncta and as shown above, the PS staining at the plasma membrane was significantly diminished in ΔWD cells (Fig 4D). Interestingly, the vesicles containing LacC2 and Rab7 in ΔWD MEFs were larger and swollen compared to WT cells. Quantitative assessments revealed a correlation between PS and Rab7 that is 4.5-fold greater in ΔWD MEFs than WT cells and indicates that more PS is found in late endosomes in cells lacking the ATG16L1 WD domain. Incubation of ΔWD cells with T0901317 markedly reduced numbers of puncta double positive for Rab7 and Lact-C2-GFP (Fig 4D). The Pearson coefficients showed that T0901317 reduced colocalization of PS with Rab7 by 3.5-fold in ΔWD cell and restored PS levels at the plasma membrane (Fig 4D). Moreover, vesicles containing cholesterol visualized using the D4H mCherry probe and Rab7 in ΔWD MEFs were also larger and swollen compared to WT cells (Fig 4E). The co-localization of D4H mCherry and Rab7 was 3.7-fold greater in ΔWD MEFs than WT cells, demonstrating that cholesterol is significantly more abundant in late endosomes of cells lacking the ATG16L1 WD domain. In summary, in cells lacking the WD domain of ATG16L1, PS and cholesterol redistribute from the plasma membrane to late endosomes containing Rab7. T0901317 treatment resulted in a markedly reduced frequency of Rab7 and D4H mCherry double-positive puncta staining in ΔWD cells (Fig 5E). Pearson coefficients suggested co-localization in vesicles to be reduced 2.8-fold by T0901317 suggesting that the drug caused loss of cholesterol from late endosomes/lysosomes in ΔWD cell. T0901317 had little effect on WT MEFs apart from a small non-statistically significant (P=0.2156) increase in D4H mCherry and Rab7 colocalization. Experiments comparing localization of cholesterol in lysosomes and early endosomes in WT and DWD cells showed partial colocalization of D4H cholesterol probe with LAMP1-positive lysosomes, but very little colocalization with EEA1-positive early endosomes (Fig 5A). Taken together, these results indicated that altered cholesterol and PS transport in ΔWD cells results in loss of cholesterol from the plasma membrane and increased cholesterol in lysosomes and late endosomes, but not in early endosomes, which are derived from membrane recycled from the plasma membrane.

**Figure 5.**
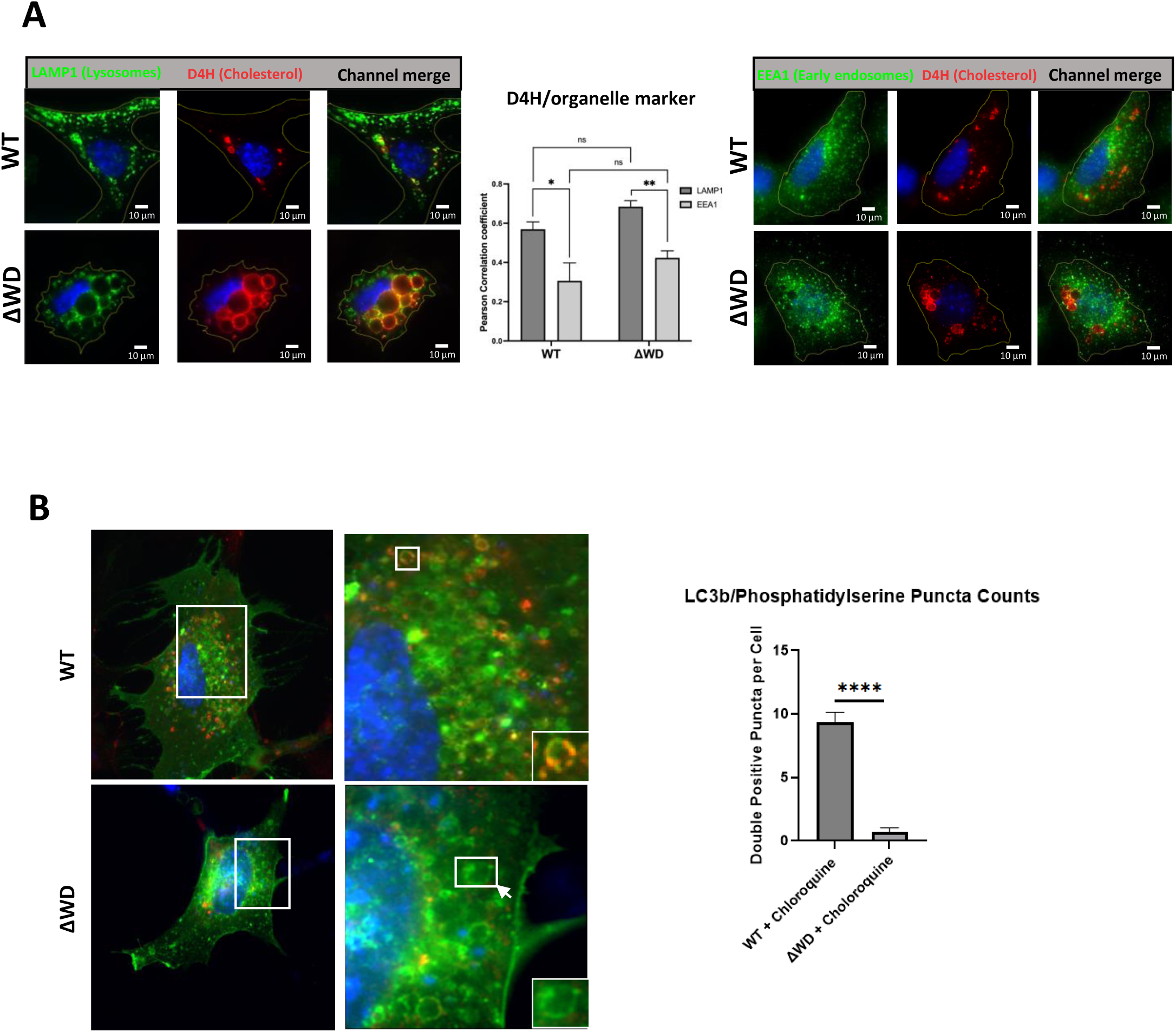
(**A**) Distribution of lysosomes and early endosomes with cholesterol in WT and ΔWD cells. Cells were transfected with D4H and immunostained with either LAMP1 or EEA1 antibodies. Pearson correlation coefficients of D4H mCherry colocalization with LAMP1 or EEA1 are presented on the graph (+SEM). Mann Whitney U test performed: ** = p<0.01, n=14. (B) WT and ΔWD MEFs transfected with Lact-C2-GFP were treated with chloroquine (50ug/ml) for 2 hours before fixing in methanol and staining with mouse anti-LC3B antibody with anti-mouse 594 secondary antibody (red). Bar chart shows quantitative counts of LC3B/Lact-C2-GFP double positive intracellular puncta using ImageJ, with graphing and statistical analysis (unpaired t-test) using GraphPad Prism (WT n=10, ΔWD n=11).

Loss of ATG5 and ATG12 also leads to cholesterol accumulation in lysosomes (22). This accumulation is independent of canonical autophagy because cholesterol distribution is not altered in cells lacking ATG3, ATG9 and ATG14 which act upstream of ATG5-ATG12. (22). This makes it possible that WD domain-dependent conjugation of LC3 to membranes regulates cholesterol and PS transport. The distribution of PS and LC3 were therefore studied in cells incubated with chloroquine to raise vacuole pH to activate CASM and LC3 conjugation to endo-lysosome membranes. In WT cells chloroquine induced the formation of swollen vesicles containing PS and many were positive for LC3. This is consistent with current models where cholesterol released from lipid droplets delivered to lysosomes by autophagy remains trapped in the lysosome because export from the vacuole through Niemann-Pick type C (NPC) transporters requires low pH (35, 36). Vesicles containing PS were also present in cells lacking the WD domain, again suggesting that PS accumulates because NPC transporters are inhibited by raised pH, but these vacuoles were negative for LC3. This suggests that WD-dependent conjugation of LC3 to vacuole membranes does not play a major role in determining PS and cholesterol distribution. This raises the possibility that the WD domain may function at other sites in cholesterol transport possibly modulating non-vesicular lipid transport mediated by lipid transfer proteins and oxysterol binding protein (OSBP)-and OSBP-related proteins (ORPs). OSBP mediates cholesterol transfer from the endoplasmic reticulum to lysosomes at membrane contact sites (37) and reverse transfer to the ER is facilitated by ORP1 and ORP5 (38).

### Cholesterol is reduced in the plasma membrane of ΔWD cells and tissues

In the next series of biochemical experiments, we used subcellular fractionation of MEF cells isolated from WT and ΔWD mice to analyze cholesterol levels in the plasma membrane. In the initial lysate, there was no significant difference in the total amount of cellular cholesterol or unesterified cholesterol between WT and ΔWD MEF cells (Fig 6A). Cells were then fractionated by density gradient centrifugation and the plasma membrane was identified by immunoblotting for β1-integrin (Fig 6B). Fractions containing the highest levels of β1 integrin across the gradient were analyzed for cholesterol content and normalized to protein concentration. There was significantly less cholesterol and unesterified cholesterol in plasma membrane of ΔWD MEFs compared to WT MEFs (Fig 6C).

**Figure 6.**
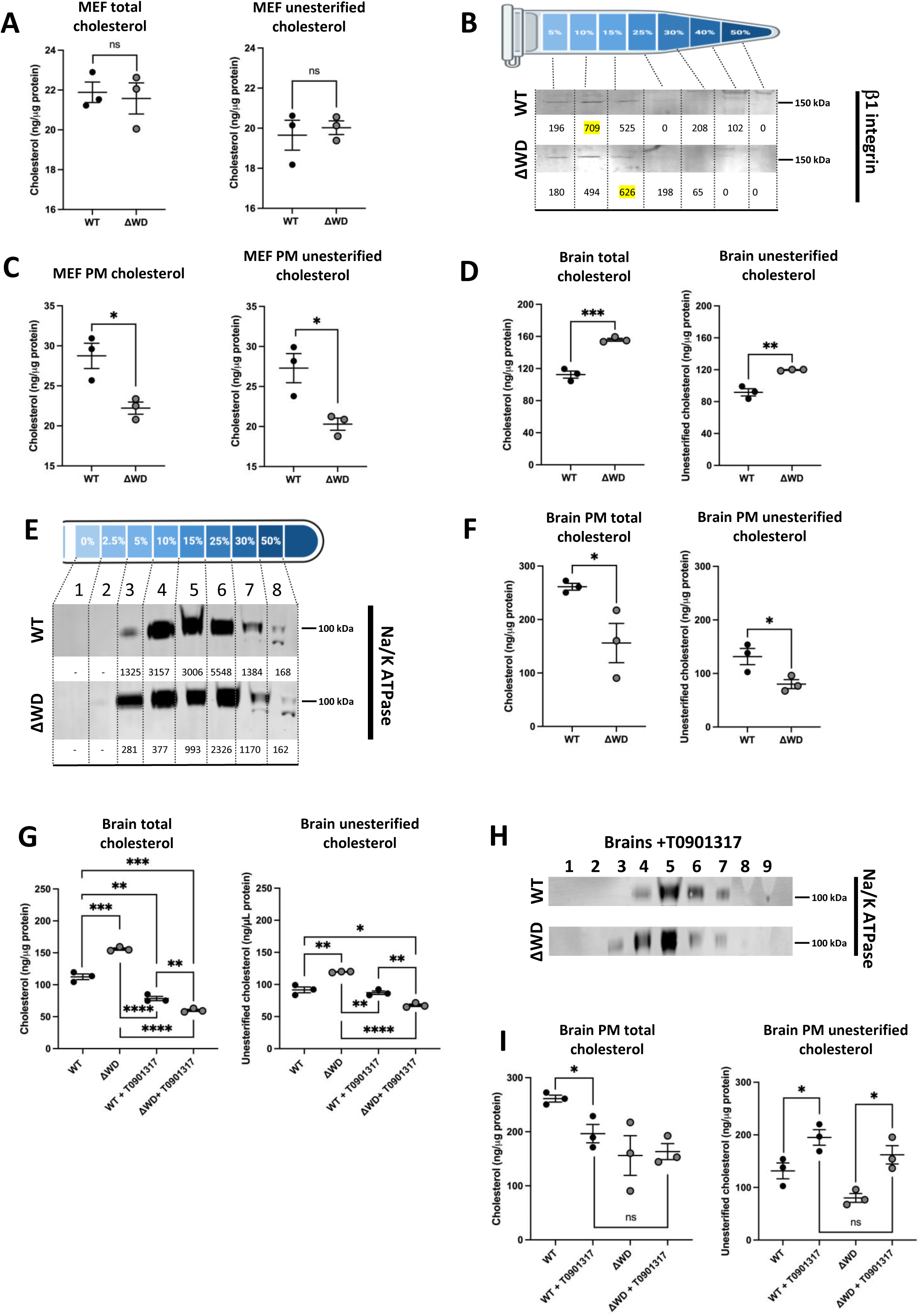
Plasma membrane in brains of mice lacking WD domain of ATG16L1 are deficient in cholesterol (**A**) Total and unesterified cholesterol was quantified from MEF lysates using the Amplex Red cholesterol assay, and normalized to the protein concentration of the WT and ΔWD samples (±SEM). Independent samples t-test performed: ns = p>0.05 N=3. (**B**) Western blot showing MEF cells following fractionation by ultracentrifugation on a 2.5-50% Nycodenz-sucrose gradient and the plasma membrane (PM) fractions identified by β1 integrin western blot, quantified on ImageJ (numbers below respective band). (**C**) Total and unesterified cholesterol was quantified from MEF PM fractions, identified in Fig 6B. Graphs report cholesterol concentration normalized to protein concentration of WT and ΔWD samples (±SEM). Independent samples t-test performed: * = p<0.05 n=3. (**D**) Total and unesterified cholesterol was quantified from brain tissue lysates. Graphs report cholesterol concentration normalized to protein concentration of WT and ΔWD samples (±SEM). Independent samples t-test performed: ** = p<0.01, *** = p<0.001 n=3. (**E**) Western blot showing brain tissue following fractionation by ultracentrifugation on a 2.5-50% step Nycodenz-sucrose gradient and the plasma membrane fractions identified by Na/K ATPase beta subunit western blot, quantified on Image J (numbers below respective band). (**F**) Total and unesterified cholesterol was quantified from brain PM fractions, identified in Fig 6E. Graphs report cholesterol concentration normalized to protein concentration of WT and ΔWD samples (±SEM). Independent samples t-test performed: * = p<0.05 n=3. (**G**). Total and unesterified cholesterol was quantified from brain tissue lysates of mice treated or untreated with T0901317 for 3 days. Graphs report cholesterol concentration normalized to protein concentration of WT and ΔWD samples (±SEM). Independent samples t-test performed: * = p<0.05, ** = p<0.01, *** = p<0.001, **** = p<0.0001 n=3. (**H**) Western blot showing brain tissue following treatment of mice with T0901317 for 3 days, fractionated on a 2.5%-50% Nycodenz-sucrose gradient and the plasma membrane fractions identified by Na/K ATPase beta subunit western blot. (**I**) Total and unesterified cholesterol was quantified from brain PM fractions of mice treated or untreated with T0901317 for 3 days. Graphs report cholesterol concentration normalized to protein concentration of WT and ΔWD samples (±SEM). Independent samples t-test performed: * = p<0.05. Figure S1. ATG16 L1 WD domains attenuated inflammatory signaling through the TLR3 pathway. (A) Expression of IFNβ1 and ISG15 mRNA in WT and ΔWD MEFs RNA (+SEM) by qPCR either untreated, or 4 hours after transfection with poly IC, or treated with TLR3 inhibitor one hour prior to transfection with poly IC. Independent samples t-test: *** = p<0.001. n=3. (B) RIG-I KO cell lines were generated by infecting WT and ΔWD MEFs with custom CRISPR gRNA lentivirus transduction particles. Knock-down of RIG-I protein expression was evaluated by western blot. WT RIG-I KO #2 and ΔWD RIG-I KO #2 cell lines were chosen for IAV challenge. (C) IFNβ1 mRNA expression (+SEM) following challenge with IAV for 120 mins in WT and ΔWD cells with inhibition of either TLR3 or RIG-I pathways. Independent samples t-test: * p<0.05 n=3.

The effect of loss of the ATG16L1 WD domain on cholesterol distribution in tissues *in vivo* was tested by analyzing membrane fractions isolated from brain tissue from ΔWD and WT mice. Interestingly, measurement of total cholesterol (Fig 6D) showed that there were significantly higher levels of cholesterol in the brains of ΔWD mice compared to litter mate controls. Analysis of cholesterol esterification revealed that this increase in total cholesterol was due predominantly to increases in unesterified cholesterol. Brain tissue from WT and ΔWD mice were homogenized and fractionated on a 2.5-50% Nycodenz-sucrose step gradient to separate plasma membrane (Fig 6E). Plasma membrane fractions were identified by presence of Na/K ATPase, and fractions with the peak levels of Na/K ATPase labelling relative to protein concentration were assayed for total and unesterified cholesterol. These results demonstrated that, despite higher levels of total cholesterol in the tissue, the concentration of plasma membrane cholesterol was reduced in ΔWD brain tissue (Fig 6F). This was consistent with the *in vitro* results obtained from ΔWD MEFs.

WT and ΔWD mice were treated for 3 days with T0901317 to promote cholesterol efflux from cells. As recorded above, the brains of untreated ΔWD mice showed greater levels of total and unesterified cholesterol compared to untreated WT controls (Fig 6G). Treatment with T0901317 significantly decreased cholesterol in WT and ΔWD brain tissue, with T0901317-treated ΔWD brain significantly lower than untreated and treated WT brain. T0901317. Treated brains from WT mice showed no difference from untreated for unesterified cholesterol levels, whereas unesterified cholesterol was significantly decreased in treated ΔWD mice to below the level of that found in treated WT mice (Fig 6G). These results suggest that the drug treatment mainly leads to the efflux of unesterified cholesterol in ΔWD mice. The brains of treated mice were fractionated on 2-5-50% Nycodenz gradient and plasma membrane fractions identified by Na/K ATPase labelling as above (Fig 6H). As shown in cell culture, WT mice contained higher levels of total and unesterified cholesterol in the plasma membrane of brain tissue compared to ΔWD mice. T0901317 treatment slightly lowered levels of total cholesterol in the plasma membrane fractions from WT brains (Fig 6I) and total cholesterol was not altered in the plasma membrane in brains of ΔWD mice treated with T0901317. Importantly, however analysis of unesterified cholesterol showed that T0901317 increased levels of unesterified cholesterol at the plasma membrane in both WT and ΔWD brain tissue. After drug treatment there was no significant difference in unesterified cholesterol levels between treated WT and ΔWD brain plasma membrane. Taken together, these data demonstrate a distinct reduction of cholesterol at the plasma membranes of ΔWD MEFs as well as within brain tissue. The cholesterol efflux promoting drug T0901317 increased levels of unesterified cholesterol in the plasma membrane of brains of both genotypes, and restored levels of unesterified cholesterol in the plasma membrane of ΔWD mice to levels seen in untreated WT controls.

## Discussion

Our study reveals a new unconventional role for the WD40 domain of ATG16L1 in the control of cholesterol distribution, plasma membrane integrity and influenza virus infection. Previous study on the ATG16L1 protein reported that the WD domain is essential in restricting lethal influenza infection (14). We add to this that absence of the WD domain increases IAV entry, nuclear delivery and replication. There is a prominent accumulation in a perinuclear compartment identified as late endosomes. and supports a previously described role for the WD domain of ATG16L1 in regulation of lipid transport (22, 23), controlling the transport of cholesterol and PS from late endosomes and lysosomes to the plasma membrane. Pharmacological treatment with a drug that upregulates the ABCA1 transporter, T0901317, restored the recruitment of both cholesterol and PS from the intracellular puncta to the plasma membrane. Image analysis identified the intracellular puncta as late endosomes, which is consistent with the finding that ATG16L1 promotes lysosome exocytosis to promote plasma membrane repair through a Rab7 late endosome pathway (22). Using sensitive biochemical analysis, there were higher cholesterol levels in the brains of mice lacking the WD domain however there was a deficiency of cholesterol and phosphatidyl serine in the plasma membrane of brain tissue. These results confirm previously published literature that ATG16L1 is involved in non-canonical pathways independent from classical autophagy. Other non-canonical roles for ATG16L1 include a protective response towards Alzheimer disease, where lack of WD domain has been shown to lead to deposition of amyloid β peptide (Aβ) through reduced Aβ receptor recycling (17).

Our results suggest that WD-dependent conjugation of LC3 to vacuole membranes does not play a major role in determining PS and cholesterol distribution. The WD domain may function at other sites in cholesterol transport possibly modulating non-vesicular lipid transport mediated by lipid transfer proteins and oxysterol binding protein (OSBP)-and OSBP-related proteins (ORPs). OSBP mediates cholesterol transfer from the endoplasmic reticulum to lysosomes at membrane contact sites. Interestingly, Rab7 is recruited to phagosomes and required for phagolysosome maturation. However, in cholesterol-loaded cells, Rab-7 is inactive with accumulation of unesterified cholesterol in endo-membranes which prevents their fusion with lysosomes (39). This may suggest that there may be an impairment of Rab 7 activation in cells lacking the WD domain, where there is an accumulation of cholesterol in late endosomes. Cells also transport lipids between organelles by non-vesicular pathways, including movement of cholesterol to the plasma membrane, a process that permits rapid membrane expansion of intracellular compartments, such as autophagosomes. Other WD-containing proteins such as ATG2A play a role in membrane expansion (39, 40). It has been well established that viral entry can be modulated by the concentration of cholesterol within the membranes of the cell-virus interaction (18); a significant presence of cholesterol within membranes orders bilayer lipids and stabilizes the fusion intermediates to promote entry (41). The use of methyl-β-cyclodextrin (MβCD) dependent depletion of cholesterol has delineated the requirements of cholesterol for specific viruses, with HIV-1 requiring cholesterol in both viral and host membranes for efficient fusion (42, 43) whereas only viral membrane cholesterol composition effected IAV fusion (44). Here we show an enhancement of viral fusion at the plasma membrane when it is comprised of less cholesterol due to the aberration of cholesterol transport through absence of the WD domain. While we cannot say definitively that this is entirely dependent on the reduced plasma membrane cholesterol, *in vitro* experiments supplementing ΔWD cells with cholesterol reduced fusion of IAV with the plasma membrane. Furthermore, treatment with U18666A, which depletes plasma membrane cholesterol, increased IAV fusion in WT cells, indicating that reduced cholesterol promotes IAV fusion. Early endosomes formed after IAV internalization from the cholesterol reduced plasma membrane may also have depleted cholesterol to promote IAV fusion. We also show that endocytosis and nuclear entry of IAV increased in cells lacking WD domain. While host membrane cholesterol MβCD depletion has been shown to not effect IAV entry (19), cholesterol depletion at the plasma membrane is reflected in early endosome membranes that promote cytoplasmic entry.

Autophagy proteins are involved in an array of immune pathways required for effective defense and normal functioning homeostasis. ATG16L1 is involved in regulating inflammatory responses, with ATG16L1 shown to suppress type I IFN signaling (45) and induce regulatory T cells (46). Specific ATG16L1 polymorphisms have been implicated in disease non-progression within chronically HIV-1 infected individuals, with distinct inflammatory and immune regulatory and responsiveness profiles observed (47). Moreover, increased cytokine signaling has been demonstrated in ATG16L1 mice possessing the T300A mutation (48), which increases susceptibility for caspase cleavage of the protein, effectively removing the WD domain, suggesting it might regulate the inflammatory response. Here we show elevated basal IFNβ transcription, with levels of ISG15 and IFIT1 expression higher in ΔWD mice following IAV challenge, facilitating an inflammatory response. ISG15 has proinflammatory properties following viral infection (49), our work supports the *in vivo* inflammatory profile. Although both RIG-I and TLR3 pathways are known to be important for IFN induction by IAV (50, 51), our experiments showed elevated signaling through the endosomal TLR3 pathway, providing evidence that the WD domain is protective against viral infection.

In conclusion, we show that removal of the WD domain from the ATG16L1 protein causes intracellular cholesterol accumulation in late endosomes/lysosomes and reduces cholesterol at the plasma membrane. The exact nature of the mechanism of WD domain in cholesterol transport requires further study. IAV infection is enhanced when this WD domain lost, and the increased viral entry is followed by a subsequent increase in IFN signaling, through the TLR3 receptor pathway and a modulation of IFN signaling, previously reported *in vivo*. Therefore, we propose that ATG16L1 not only protects cells from virus entry, but also the elevated downstream cytokine responses to infection, which in mice can progress to severe cytokine release syndrome and death.

## Methods

### Virus propagation

Influenza virus A PR8 is a mouse-adapted H1N1 strain originally derived from a human isolate. X31 is a mouse adapted H3N2 strain with the 6 internal genes of PR8 and the HA and NA derived genes from A/Aichi/2/1968. They were propagated in the allantoic cavity of 9-day-old embryonated chicken eggs at 35°C for 72 h (52) and titer determined using plaque assay on Madin Derby Canine Kidney cells (MDCK) (53).

### Reagents

The generation of ΔWD mice (*Atg16L1ΔWD/ΔWD*) has been described previously (7). Mouse Embryonic Fibroblasts (MEFs) were procured from mice at embryonic day 13.5 and cultured in Dulbecco’s Modified Eagles Medium (DMEM) with GlutaMAX (ThermoFisher#10567014) supplemented with 1% penicillin/streptomycin, kanamycin and 10% Fetal bovine serum (Gibco, 10082147). Bone Marrow Derived Macrophages (BMDMs) were generated from bone marrow isolated from femur and tibia flushed with RPMI 1640.

Macrophages were generated by culturing adherent cells in RPMI 1640 (ThermoFisher#11835030) containing 10% FCS and CSF1/M-CSF (R&D systems, 30 ng/mL) for 6 days. Filipin III was from Sigma (#F4767). TRIzol was from ThermoFisher (#15596026); WGA 647 was from Invitrogen (W32466 5 µg/mL); cholesterol was from Sigma Aldrich C8667 80 µM; 25-hydroxycholesterol was from Sigma Aldrich (H1015 5 µM) TLR3 inhibitor was from Sigma-Aldrich (#614310); poly I:C was from Sigma-Aldrich (#P1530). RIG-I knock-out (KO) cell lines were generated by infecting WT and ΔWD MEFs with custom CRISPR gRNA lentivirus transduction particles (Sigma Aldrich MMPD0000132807 and MMPD000132808) in Opti-Minimal Essential Medium (Opti-MEM) supplemented with 16 µg/mL hexadimethrine bromide. Selection was performed using 10 µg/mL puromycin and knock-out validated by western blotting using anti-RIG-I/DDX58 (Abcam ab180675). The Lact-C2-GFP plasmid was from Addgene (#22852) TLR9 ligand OD1585 was from InvivoGen

### Lung slice culture

The ex vivo lung tissue model has been previously described (54). Briefly, WT and ΔWD mouse lungs were harvested and airways were inflated with 1% low melting point agarose in DMEM F12 (Thermofisher#10565018) in Hanks Balanced Salt Solution (HBSS, ThermoFisher#14025092) and cast in blocks of 2% agarose in HBSS for sectioning with a vibrating microtome. Slices were cut at 300 µm and incubated in DMEM F12 + penicillin/streptomycin + kanamycin, overnight at 37°C 5% CO2 and infected with IAV as indicated.

### Drug treatment

MEFs were treated with 10μM/mL LXR agonist T0901317 (Sigma Aldrich#T0901317) diluted in complete media for 48 hours prior to harvesting for ultracentrifugation. For mouse treatment, T0901317 was administered for 3 days prior to sacrifice at concentrations of 25 mg per kilogram of body weight. In tissue culture, U18666A (Cayman Chemical#10009085) was used at 3 µg/mL for 24 hours. Cholesterol and 25-hydroxy cholesterol (25HC) were prepared in ethanol and used at 80 µM and 5 µM respectively. The TLR3/dsRNA complex inhibitor C18H13ClFNO3S (Sigma) blocks dsRNA binding to TLR3 and was added one hour prior to infection or transfection. Chloroquine (50ug/ml) was added to cells for 2 hours before fixing.

### IAV entry assays by acid mediated bypass

We adapted methods previously published methods to force fusion of the IAV membrane to the plasma membrane (24, 55). MEFs were seeded onto coverslips (NP entry assay) or 6-well plates (IFNβ assay) and grown to 75% confluency or 100% confluency respectively. Cells were infected with IAV X31 at an MOI of 100 for the NP entry assay and 10 for the IFNβ assay. Virus was bound to MEFs for 1 hour at 4°C, unbound virus was removed via cold infection medium (DMEM, 50 mM HEPES, pH 6.8, 0.2% BSA).

Cells were incubated for 2 min in FUSION medium (DMEM, 50mM citric buffer adjusted to pH 5.0) before being cooled, washed in cold infection medium and incubated at 37°C in STOP medium (DMEM, 50mM HEPES, 20mM NH4Cl, pH 7.4) for the appropriate time points. A control condition was also performed using pH 7.0 medium instead of the FUSION medium.

### Endocytosis Assay

High-resolution analysis of IAV entry to cells via endocytosis was measured using a previously published protocol (24, 55). IAV X31 (MOI: 10) was diluted in infection medium (DMEM, 50 mM HEPES, pH 6.8, 0.2% BSA) and used to infect MEFs at 4°C for 1 hour. Pre-treatment of cells with Dynasore (Sima-Aldrich# 324410, 80 µM) for 30 min was used as a control. Unbound virus was removed with ice cold infection medium before bound virus was internalized in warm media at 37°C for a 30 min incubation. Cells were then fixed in 4% paraformaldehyde (PFA) after which the plasma membranes were stained (WGA 647) before cells were blocked for 30 min (1% BSA, 5% FCS, 1 x PBS). External influenza HA epitopes were immunostained with the primary antibody PINDA (made by Y. Yamauchi lab) (1:500) in a blocking solution and incubated overnight at 4°C. PINDA was then stained with anti-rabbit IgG-Alexa Fluor® 594 (Abcam) (1:1000) in blocking solution for 1 hour at room temperature. Cells were fixed again in 4% PFA and permeabilized in 0.1% Triton™X100, 1% BSA, 5% FCS, 1 x PBS. Cells were incubated with HA1 antibody (1:100), specific for the internalized HA1 epitope. Cells were washed and stained with the anti-mouse IgG Alexa Fluor® 488 (1:1000). Nuclei were stained with DAPI (1:5000) before coverslips were mounted and viewed using a Zeiss Axio Imager 2 for representative images. For automated image acquisition a 20X lens was used on a Zeiss LSM laser scanning confocal microscope, measuring over 3 independent cover slips. Images were analyzed using the CellProfiler program, with a spot detection algorithm employed for detection of puncta and categorizing them, which is detailed in (24, 55), counting 880 WT cells and 419 ΔWD. An in-house Python script was used to remove cells with no virus puncta, resulting in 837 WT cells and 417 ΔWD being used in analysis.

### Nuclear entry assay

IAV was used to infect primary cells (MOI: 4) and diluted in OptiMEM. (Thermo Fisher 31985070) WT and ΔWD MEFs were seeded onto on coverslips. Virus was bound to cells at 4°C for 1 hour, then allowed to internalize through incubation at 37°C. Cells were fixed in 4% PFA at the following time points: 2, 3, 4 and 5 hpi; permeabilized with 0.1% Triton x100; blocked in 0.1M glycine and 2% BSA solutions; and immunostained with anti-IAV NP (Abcam ab20343) in 2% BSA blocking solution overnight at 4°C. Cells were then stained with IgG Alexa Fluor® 488 (Abcam ab150105) in 2% BSA blocking solution for 2 hours at room temperature. Nuclei were stained with DAPI (Invitrogen #D1036) for 10 min before being mounted and viewed using a Zeiss Axio Imager 2 microscope.

### Quantitative IAV and cytokine transcription

Tissues were frozen in liquid nitrogen and homogenized using a TissueLyser with TRIzol. Tissue culture cells were washed twice using PBS before administration of TRIzol and cell scraper/23G needle homogenization. With both tissue and cells TRIzol–chloroform extractions were further purified by RNeasy MinElute Cleanup Kit according to manufacturer’s instructions. RNA was analyzed by quantitative PCR (qPCR) using SYBR Green/7500 Standard Real-Time PCR System and Qiagen primer sets. IAV M forward primer 5’ GACCRATCCTGTCACCTCTGAC 3’ and M Reverse: 5’ AGGGCATTYTGGACAAAKCGTCTA 3’ (56) Relative amounts of mRNA expression was normalized to 18S rRNA.

### Subcellular fractionation of Mouse Embryonic Fibroblast

WT control and ΔWD MEFs were cultured until confluent, harvested by scaping and homogenate pelleted at (14,000 xg, 5 mins, 4°C). The pellet was homogenized on ice by passaging through a 23G needle and lysed cells pelleted at (100 xg, 10 minutes, 4°C) with post nuclear supernatant (PNS) collected. MEF homogenates were fractionated using a discontinuous step gradient (5-50%) of sucrose-Nycodenz across a total volume of 1.4 ml. Tubes were loaded into a Beckman TLS-55 ultracentrifugation rotor. Samples were separated by ultracentrifugation at (61,725 xg, 3 hours, 4°C). Afterwards, 7 x 200µl fractions were collected with protein concentration in PNS and subcellular fractions was quantified using a bicinchoninic acid (BCA) assay.

### Subcellular fractionation of Mouse Brain

WT and ΔWD mice were sacrificed and brains extracted, weighed, and minced. Tissue was collected and pelleted by centrifugation. The pellet was resuspended in ice cold hypotonic buffer (dH20, 10mM Tris-HCl, and 1mM EDTA), and incubated on ice for 1 minute. Hypotonic buffer was removed through centrifugation before resuspension in 2ml of buffered sucrose solution and homogenization using a Dounce homogenizer. Lysates were sonicated on ice for 30 seconds twice before pelleting to remove nuclei, with the PNS fraction collected. Subcellular fractionation of mouse brains was performed on a 32 mL discontinuous (0-30%) Nycodenz gradient in a Beckman SW32 Ti ultracentrifuge rotor. Samples were separated by ultracentrifugation (59,333 xg, 3 hours, 4°C). Afterwards, 8 x 3.5ml fractions were collected and stored at −20°C. Protein concentration in PNS and subcellular fractions was quantified using BCA.

### Protein Expression

MEF plasma membrane fractions were detected using polyclonal β1-Integrin staining (Uli Mayer, University of East Anglia). Brain plasma membrane fractions were detected with a Na/K ATPase monoclonal antibody. For plasma membrane isolation, protein bands were quantified through automated densitometry performed using Odyssey CLx software. Densitometry readings were then standardized against fraction protein concentration, determined via BCA, with fractions containing the highest concentration of plasma membrane protein denoted as the plasma membrane peak fractions and assayed for cholesterol concentration.

### Biochemical analysis of Cholesterol

Cholesterol concentrations were measured using the Amplex® Red cholesterol assay kit (Invitrogen) according to the manufacturer’s instructions. Resorufin fluorescence was measured on a SpectraMax M2microplate reader (excitation: 545 nm, emission: 590 nm). Cholesterol quantifications were then normalized to protein concentration of the sample as determined by BCA.

### Lipid localization

Cholesterol was localized using either D4H mCherry probe or filipin. D4H mCherry plasmid was kindly provided by Gregory D. Fairn (33). Lact-C2-GFP and IFITM3-Myc plasmids were purchased from Addgene. Plasmids were purified using a NucleoBond® Xtra Midi prep kit. D4H mCherry, Lact-C2-GFP, and IFITM3-Myc were transfected into WT and ΔWD MEFs in a 24-well plate using Lipofectamine 3000 reagent. Cells were then fixed using 4% PFA for 10 minutes; permeabilized in 0.1% Triton X100 for 10 minutes; blocked in 0.1M glycine and 0.5% BSA solutions for 20 and 30 minutes respectively; and stained with either anti-Rab7 (Late endosomes), anti-LAMP1 (lysosomes) or anti-EEA1 (Early endosomes) at 4°C overnight. Staining with Alexa Fluor® 488 or 594 was performed before mounting and imagine using either a Zeiss M2 Imager or a Zeiss LSM980-Airyscan confocal microscope. For filipin staining of tissue: tissues were frozen using optimal cutting temperature compound (OCT) in liquid nitrogen chilled isopentane. Specimens were cut (10 µm) and mounted onto a microscope slide, fixed in 4% PFA and stained with 0.5 mg/mL filipin III in PBS + 1% BSA for 2 hours at RT. Lung slices were also stained with anti-rab7. For filipin staining of MEFs: cells were fixed with 4% PFA and stained in 0.5 mg/mL filipin III in PBS + 1% BSA for 2 hours at room temperature. Coverslips were mounted and viewed using a Zeiss LSM980-Airyscan confocal microscope with the 405 nm laser.

### Statistics

For comparative analysis between two groups, Mann Whitney U tests and independent samples t-tests were carried out where appropriate. For imaging, background staining exclusion and Pearson’s correlation coefficients determined using Imaris imaging colocalization software and ImageJ. CellProfiler and Python were used in the acquisition and processing of the endocytosis assay data.

## Abbreviations

IAV: influenza virus A
LAP: LC3-associated phagocytosis
WD: WD40 -repeat-containing C-terminal domain
LDL: low density lipoprotein
AD: Alzheimer’s disease
ISG: interferon-stimulated gene
IFN: interferon
PS: phosphatidylserine
CCD: coil-coli domain
WT: wild type
BMDM: bone marrow derived macrophages
LAP: LC3-associated phagocytosis
CASM: conjugation of ATG8 to single membranes
LANDO: LC3-associated endocytosis

## Acknowledgements

We thank Gregory Fairn, Dept of Biochemistry University of Toronto, Toronto, Ontario for the D4H-mcherry plasmid, James McColl (Faculty of Science, UEA) for help with confocal imaging and Naiara Beraza (Quadram Institute) for T0901317 treatments.

