## Supplementary figures and images for "ATG16L1 WD domain regulates lipid trafficking to maintain plasma membrane integrity to limit influenza virus infection"

### SupplementalFigure 1

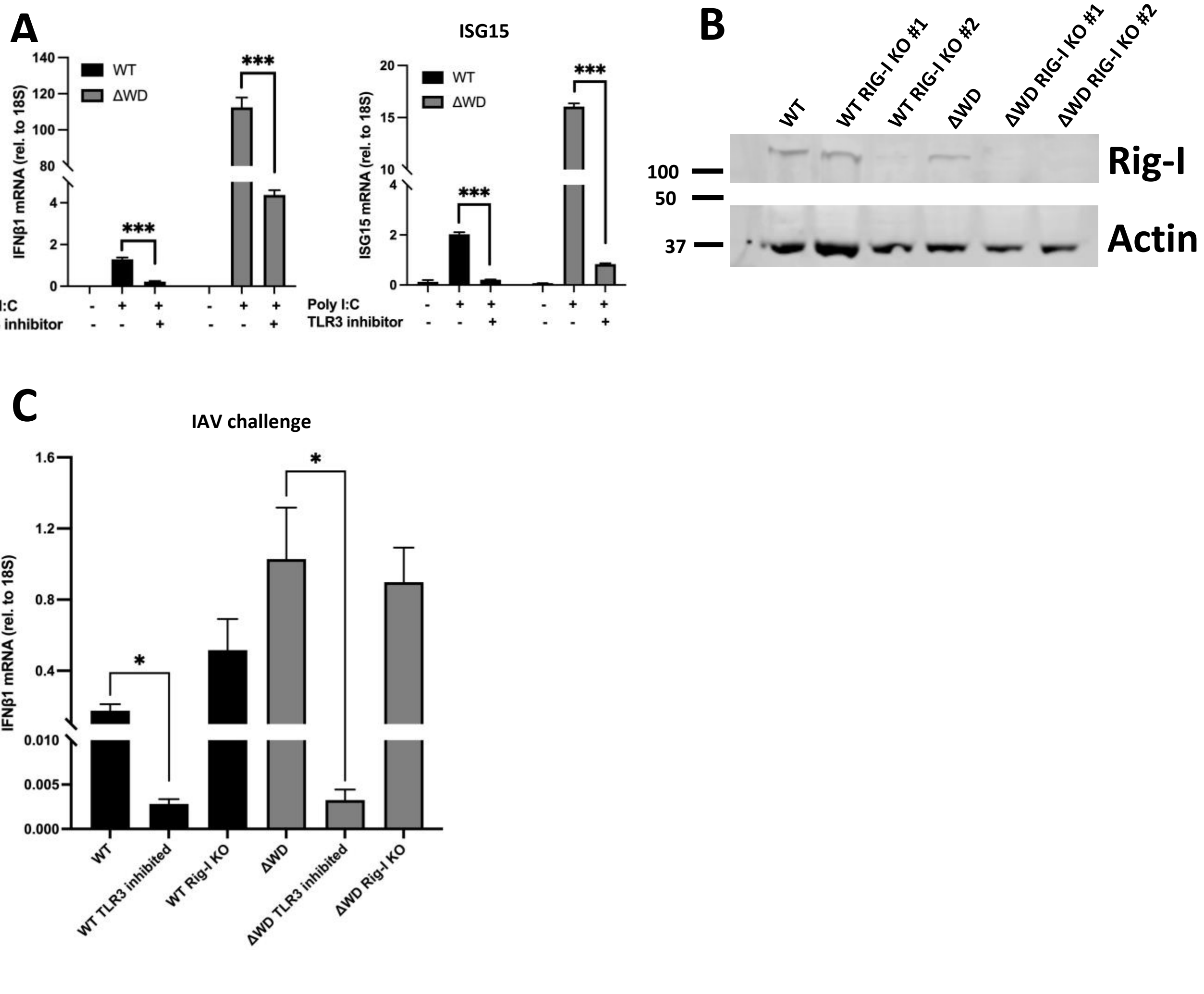
